# Evaluation of the cross reactivity of neutralising antibody response in vaccinated human and convalescent hamster sera against SARS-CoV-2 variants up to and including JN.1 using an authentic virus neutralisation assay

**DOI:** 10.1101/2023.10.21.563398

**Authors:** Naomi S. Coombes, Kevin R. Bewley, Yann Le Duff, Nassim Alami-Rahmouni, Kathryn A. Ryan, Sarah Kempster, Deborah Ferguson, Elizabeth R. Davies, Thomas M. Weldon, Eleanor S. Cross, Lauren Smith, Conner Norris, Karly Rai Rogers-Broadway, Kuiama Lewandowski, Samantha Treagus, Isobel Everall, Steven T. Pullan, Bassam Hallis, Sue Charlton, Yper Hall, Simon G. P. Funnell

## Abstract

New vaccines, therapeutics and immunity elicited by natural infection create evolutionary pressure on SARS-CoV-2 to evolve and adapt to evade vaccine-induced and infection-elicited immunity. Vaccine and therapeutics developers thus find themselves in an “arms race” with the virus. The ongoing assessment of emerging SARS-CoV-2 variants remains essential as the global community transitions from an emergency response to a long-term management plan. Here, we describe how an authentic virus neutralisation assay using low passage clinical virus isolates has been employed to monitor resistance of emerging virus variants to neutralising antibodies from humans and experimentally infected hamsters. Sera and plasma from people who received three doses of a vaccine as well as those who received a bivalent booster were assessed against SARS-CoV-2 variants, up to and including JN.1. Contemporary or recent virus variants showed substantial resistance to neutralisation by antibodies from those who had received three doses of an ancestral vaccine but were still effectively neutralised by antibodies from individuals who had received a bivalent booster (ancestral/BA.1). In our recent studies, however, the JN.1 VOI was found to be significantly more resistant to neutralisation by antibodies from those who had received the ancestral/BA.1 bivalent boost. Convalescent sera from hamsters that had been experimentally infected with one of seven virus variants (ancestral, BA.1, BA.4, BA.5.2.1, XBB.1.5, XBB.1.16, XBB.2.3) were also tested here. The recent contemporary variant, BA.2.86, was effectively neutralised by sera from hamsters infected with XBB.1.5 and XBB.1.16 but it was not neutralised by sera from those infected with BA.5.2.1. These data support the recommendations given by the WHO that a new vaccine was required and should consist of an XBB sub-lineage antigen.

## Introduction

Coronavirus 2019 disease (COVID-19) caused by SARS-CoV-2 continues to cause significant disruption to healthcare systems and economies worldwide having caused over 774 million cases and 7.01 million deaths since its emergence in December 2019 (as of January 2024) [1]. The WHO downgraded the COVID-19 pandemic from a public health emergency of international concern (PHEIC) on 5 May 2023, this marked a transition from emergency response to long-term management [2]. However, the virus continues to evolve in response to changing population immunity leading to the regular identification of new viral variants. Sequences of emerging variants are continuously monitored and assigned as new sub-lineages by the Phylogenetic Assignment of Named Global Outbreak Lineages (PANGOLIN) tool. At the time of writing there are 3,569 designated SARS-CoV-2 lineages with five of these having been classified as variants of concern (VOC) by the WHO Technical Advisory Group on Virus Evolution: Alpha (B.1.1.7), Beta (B.1.351), Delta (B.1.617.2), Gamma (P.1) and Omicron (B.1.1.529) [3–5].

A global effort to design and manufacture COVID-19 vaccines began as soon as the viral genome was published and the first COVID-19 vaccine was deployed in the UK on 8^th^ December 2020 (Comirnaty) [6]. This was followed by Vaxzevria on 30^th^ December 2020 and Spikevax on 8^th^ January 2021 [7, 8]. These vaccines were the most widely used worldwide for the initial course of three immunisations and all were based on the ancestral (clade A) virus spike glycoprotein. In the first year of vaccines being available it has been predicted that vaccination prevented as many as 14.4 million deaths due to COVID-19 [9]. The emergence of novel SARS-CoV-2 variants brought with it the concern that ancestral-based vaccines would no longer provide sufficient protection to vulnerable people in at-risk groups. Indeed, our data utilising the gold-standard authentic virus neutralisation assay indicated that the latest variants were becoming increasingly resistant to ancestral-based immune responses [10–12]. Bivalent vaccines were developed in response to this concern and have been designed to not only target the ancestral strain but also Omicron sub-lineage spike glycoproteins. These were approved for use in the UK (BA.1/ancestral bivalent) for at-risk groups only in the Autumn of 2022, and the US (BA.5/ancestral bivalent) for anyone aged 18 or over in August 2022 [13, 14]. Restricting vaccination to only at-risk groups in the UK and low vaccine uptake in the US, where only 34% adults received a fourth booster, means that most of the population who have received a vaccination are still immunised with an ancestral spike-based vaccine [15]. However, since the relaxing of restrictions worldwide, many people will also have received some degree of natural immunity due to infection with one or more of the circulating variants. The most recent vaccines to be approved elicit immunity against contemporary XBB recombinants, the latest to gain authorisation consists of an XBB.1.5 component [16–18].

The recent emergence of the BA.2.86 variant and its subvariants such as JN.1 has highlighted the value of sustained sequence monitoring due to the sudden appearance of a virus with vastly different spike sequence compared to other circulating variants or those represented in the latest vaccines. As part of a CEPI-funded “Agility” project, we have been monitoring and performing risk assessments of SARS-CoV-2 variants since November 2020. We previously presented our analysis of 20 different variants using early pandemic human convalescent and triple-vaccinated plasma and serum panels as well as international standards [12]. Here we describe our latest data generated using serum or plasma collected from a panel of individuals who have received up to four doses of a COVID-19 vaccine using an authentic virus neutralisation assay. These assays have been performed using well characterised, low passage SARS-CoV-2 viruses derived from clinical isolates including the BA.2.86 and JN.1 variants. We also present neutralisation data generated using the sera of experimentally infected hamsters which were convalescent from either the ancestral virus (pre-D614G), BA.1, BA.4, BA.5.2.1, XBB.1.5, XBB.1.16 or XBB.2.3 (mono-variant convalescent). We have also compared the antigenic maps of these two data sets.

## Results

Through monitoring of the currently circulating SARS-CoV-2 variants, we identified and sourced nasopharyngeal swabs from sequence-confirmed patients through our clinical networks and Pillar 1 testing (primarily positive tests conducted in primary healthcare settings) in England. We then used an authentic virus FRNT using low passage SARS-CoV-2 variants of clinical origin to perform risk assessments with regards to cross-reactivity of humoral immunity. SISPA-Illumina sequencing was performed on stocks prior to use in the FRNT to confirm the lineage and check for cell culture-induced mutations. We note that one of the BA.2.86 variants (BA.2.86 – A) described here contains an unexpected amino acid change within the furin cleavage site (FCS) (S:R682W). However, work published by others indicates that changes in this position are unlikely to alter the neutralisation profile, though this mutation makes this stock unsuitable for *in vivo* studies [19–22]. Neutralisation assays were therefore performed with this virus whilst efforts continued to generate a bank of BA.2.86 with a wild-type FCS (BA.2.86 – B). The variants used in this study, and their sources, are detailed in Table 1.

**Table 1.** SARS-CoV-2 variants used in this study.

| **WHO Designation** | **Pangolin Lineage** | **GISAID Clade/Lineage** | **Nextstrain Clade** | **Date Assigned** | **GISAID ID of Isolation Swab (Where Known) and/or Source** |
| --- | --- | --- | --- | --- | --- |
| Ancestral virus | B | O | 19A | Jan 2020 | EPI_ISL_406844 |
| Alpha | B.1.1.7 | GRY | 20I (V1) | Dec 2020 | EPI_ISL_683466 |
| Beta | B.1.351 | GH/501Y.V2 | 20H (V2) | Dec 2020 | EPI_ISL_770441 |
| Delta | B.1.617.2 | G/452R.V3 | 21A | May 2021 | EPI_ISL_2742236 |
| Zeta–FioCruz * | P.2 | GR/484K.V2 | 20B/S.484K | Mar 2021 | FioCruz, Brazil |
| Zeta–BEI * | P.2 | GR/484K.V2 | 20B/S.484K | Mar 2021 | BEI (NR-55439) |
| XE | XE | GRA | Recombinant | Mar 2022 | EPI_ISL_11586931 |
| XF | XF | GRA | Recombinant | Mar 2022 | EPI_ISL_10458256 |
| Omicron | BA.1 | GRA | 21K | Dec 2021 | EPI_ISL_7400555 |
| Omicron | BA.1.1 | GRA | 21K | Jan 2022 | EPI_ISL_8165999 |
| Omicron | BA.2 | GRA | 21L | Dec 2021 | Not available |
| Omicron | BA.2.12.1 | GRA | 22C | Apr 2022 | Gavin Screaton (University of Oxford) |
| Omicron | BA.2.75.3 | GRA | 22D | Jun 2022 | EPI_ISL_13882158 |
| Omicron | BA.2.75.2 | GRA | 22D | Aug 2022 | Not available |
| Omicron | BA.4 | GRA | 22A | Apr 2022 | EPI_ISL_13157810 |
| Omicron | BA.4.6 | GRA | 22A | Jul 2022 | Not available |
| Omicron | BA.5.2.1 | GRA | 22B | Apr 2022 | EPI_ISL_12810908 |
| Omicron | BQ.1.22 | GRA | 22E | Sep 2022 | EPI_ISL_15581064 |
| XBB.1.1 | XBB.1.1 | GRA | 22F | Oct 2022 | EPI_ISL_15682231 |
| Omicron | CH.1.1 | GRA | 23C | Oct 2022 | EPI-ISL-16615644 |
| XBB.1.5 | XBB.1.5 | GRA | 23A | Jan 2023 | EPI-ISL-16587654 |
| XBB.1.16 | XBB.1.16 | GRA | 23B | Mar 2023 | EPI_ISL_17666853 |
| XBB.2.3 | XBB.2.3 | GRA | 23E | Jan 2023 | Not available |
| EG.5.1.1 | EG.5.1.1 | GRA | 23F | Jun 2023 | EPI_ISL_18355898 |
| Omicron* | BA.2.86 (A) | GRA | 21L | Aug 2023 | EPI_ISL_18111770 |
| Omicron* | BA.2.86 (B) | GRA | 21L | Aug 2023 | EPI_ISL_18274087 |
| Omicron | JN.1 | GRA | 21L | Sep 2023 | EPI_ISL_18521510 |

\* For Zeta and BA.2.86, assays were conducted with stocks derived from viruses of different sources.

Our previously published data indicated that immunity derived from neutralising antibodies was becoming less effective for those who had received three doses of a first-generation COVID-19 vaccine against the currently circulating variants [12]. Whilst a newer panel of sera was acquired, we continued our assessment of triple vaccinated individuals who had received an original ancestral based vaccine. This panel had been tested against ten SARS-CoV-2 variants (including ancestral) at the point of publishing our previous paper. We have subsequently tested this panel against a further six variants (Zeta – FC, Zeta – BEI, BA.4.6, BA.2.75.2, BQ.1.22 and XBB.1.1), as shown in Figure 1.

**Figure 1.**
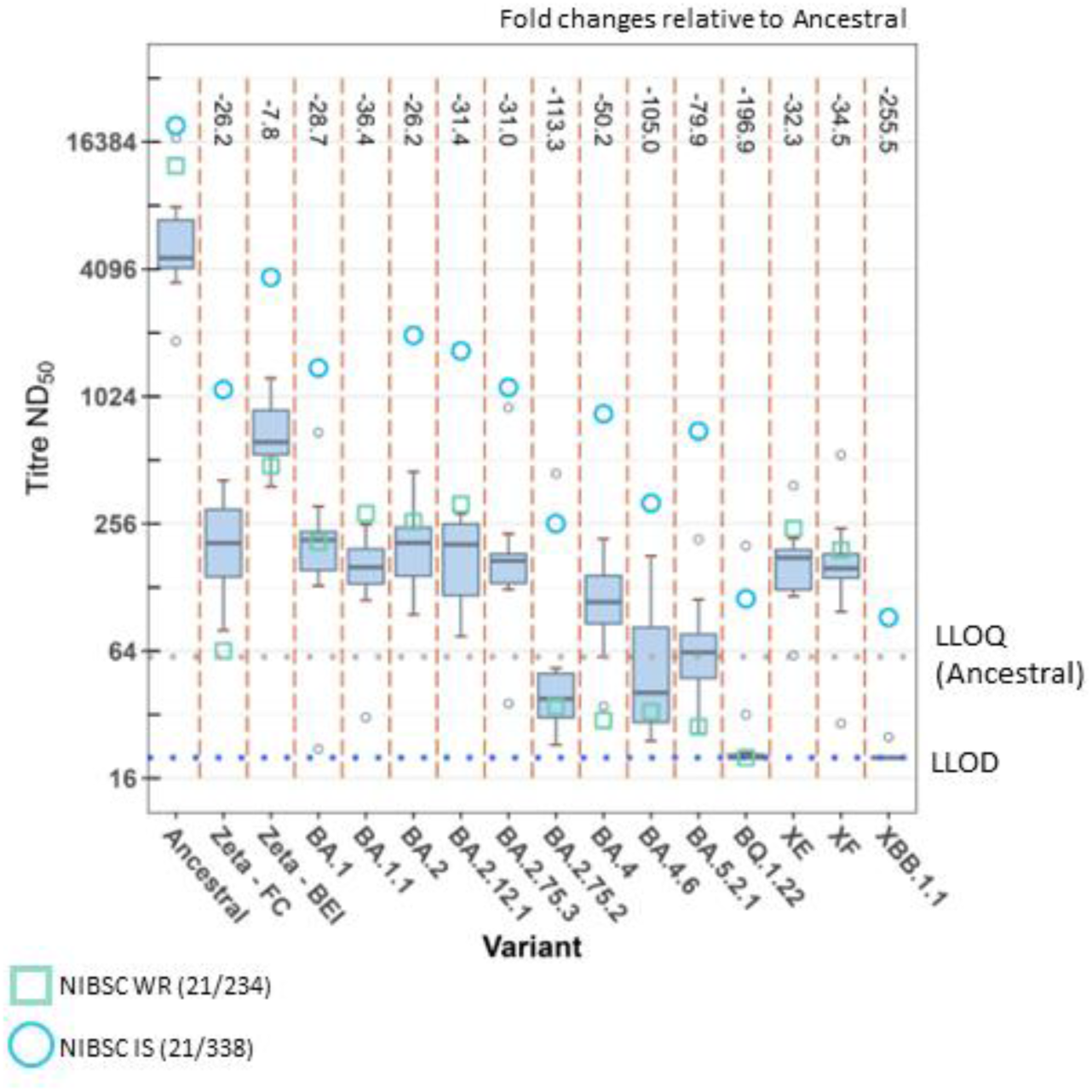
Neutralising antibody titres of a panel of triple vaccinated human participant sera determined by Focus Reduction Neutralisation Test (FRNT) against authentic SARS-CoV-2 Variants. A panel of ten sera from triple vaccinated human volunteers were assessed by FRNT with 16 authentic SARS-CoV-2 variants. Data are presented as median and interquartile ranges of titre ND_50_, titres of panel members falling outside of these ranges are plotted as open small circles. Fold-changes relative to ancestral virus were calculated by regression analysis using a mixed-effect model and are presented where the difference is statistically significant (p<0.05, two-way ANOVA with Tukey’s *post hoc* test). The Lower Limit of Detection (LLOD) of 1/20 is indicated by the blue dotted line. The Lower Limit of Quantification (LLOQ) of 1/58 as determined for the ancestral assay, is indicated by the grey dotted line. Neutralisation titres of the standards; NIBSC WR 21/234 and 21/338, are presented as a green box and blue circle, respectively.

Our previous report demonstrated a significant difference in ND_50_ titres of our pre-Alpha convalescent panel when tested against two different Zeta isolates. This was notable due to most other published data for this variant demonstrating comparable neutralisation titres to those against Delta. However, another group demonstrated a similarly large resistance to neutralisation of Zeta with a panel of early convalescent sera but not a panel of vaccinee sera [23]. We therefore sought to investigate whether the same pattern would be observed with our triple vaccinee panel (Figure 1). Geometric mean ND_50_ neutralisation titres (GM) of this panel against Zeta – FC and Zeta – BEI were 199 and 671 corresponding to a 26.2-fold and 7.8-fold reduction relative to ancestral, respectively. The difference between the titres measured against the two Zetas was significant (fold-change = 3.4; ME-ANOVA; p < 0.05). Of the four remaining additional variants that were tested against this panel since our previous paper, BA.4.6 and BA.2.75.2 resulted in GM titres above the lower limit of detection of the assay (LLOD), thus permitting more accurate estimates of fold-reduction relative to ancestral virus. The fold-reduction for BA.4.6 relative to ancestral was statistically significant at 105. When compared to its close parent lineage, BA.4, there was also a significant fold-reduction of 2.1. Against BA.2.75.2 the triple-vaccinated panel gave a GM titre of 46.1 which equated to a 113-fold reduction relative to ancestral and 3.7-fold reduction relative to a closely related variant, BA.2.75.3. Two of the most recent variants (BQ.1.22 and XBB.1.1) were substantially resistant to neutralisation to sera from triple-vaccinated individuals with most titres falling below the LLOD. Sera which fall below the LLOD are assigned the value of the lowest dilution used in the assay (1/20), thus fold-reduction calculations do not provide accurate estimates. The lower limit of quantification (LLOQ) for the ancestral virus assay (previously determined as 58) is also displayed [11]. Although the LLOQ was not determined for subsequent variants, the precision of the assay is likely to be lower for titres falling below this value. However, the estimated fold-reductions relative to ancestral virus for BQ.1.22 and XBB.1.1 were estimated at 197 and 226-fold.

In addition, to assessing the sera from individuals of the panel, the neutralisation titres of the NIBSC working reagents 21/234 and/or International Standard 21/338 were also assessed in every assay. These are displayed in Figure 1 as green squares or blue circles, respectively. As described previously, these are pools of pre-Alpha convalescent plasma (21/234) or individuals who were both vaccinated and convalescent following an Alpha, Beta or Delta infection (21/338). These standards are typically more potent at neutralising the virus compared to the panel. However, it was noted that for BA.2.75.2, BA.4, BA.4.6 and BA.5.2.1, the 21/234 standard was yielding ND_50_ values comparable to those from the lowest responders of our panel and was completely unable to neutralise BQ.1.22. The newest 21/338 standard retains good reactivity and ability to neutralise all variants assessed herein.

To continue to assess humoral immunity to currently circulating SARS-CoV-2 variants, a selection of participants who had received a fourth (booster) vaccine were sought. A panel of 22 individuals was assembled from donations given by eight MHRA staff members and 14 UKHSA staff members who had received three doses of a first-generation COVID-19 vaccine followed by a booster with a bivalent (BA.1/ancestral) vaccine. Samples were collected a median of 78 days after the most recent vaccination. Among the 22 individuals, 19 had a known previous SARS-CoV-2 infection between February 2021 and December 2022.

This four-dose panel was first assessed against the ancestral virus which resulted in the highest neutralisation titres with a GM of 12,400 (Figure 2). Next, we assessed the titres of this panel against one of the earlier Omicron sub lineages, BA.4. The GM neutralisation titres of the panel against BA.4 was 624.0 which equated to a 19.8-fold reduction relative to ancestral virus. To date this panel has been tested against a further nine variants. One of these variants with a large resistance to neutralising antibodies was CH.1.1 with a GM titre for the panel of 68.7 and fold-reduction of 180. Another Omicron sub lineage, BQ.1.22, also displayed notable resistance to neutralisation by the four-dose panel which generated a GM titre of 102.0 corresponding to a 121.9-fold-reduction relative to ancestral virus. The recombinants, XBB.1.1, XBB.1.5, XBB.1.16, XBB.2.3 and EG.5.1.1 all gave comparable GM titres between 82.9 and 137.0. Similarly, the recently emerged Omicron sub-lineage, BA.2.86, was neutralised equally well by those who have had received four doses of a COVID-19 vaccine. Here, we tested two BA.2.86 isolates with or without an FCS substitution (S:R682W), BA.2.86 – A and BA.2.86 – B, respectively. The GM titres of the panel against each virus was 99.3 and 97.0, equating to a 124.7 and 127.6-fold reduction relative to ancestral, respectively. There was no significant difference between BA.2.86 – A and BA.2.86 – B (p=1.00). The variant tested here with the largest resistance to neutralisation was the Omicron sub lineage, JN.1. The GM titre for the panel against JN.1 was observed to be 28.9 which represents a 428.9-fold reduction relative to the ancestral virus and 3.4 lower fold-reduction relative to BA.2.86 (A and B, p<0.05).

**Figure 2.**
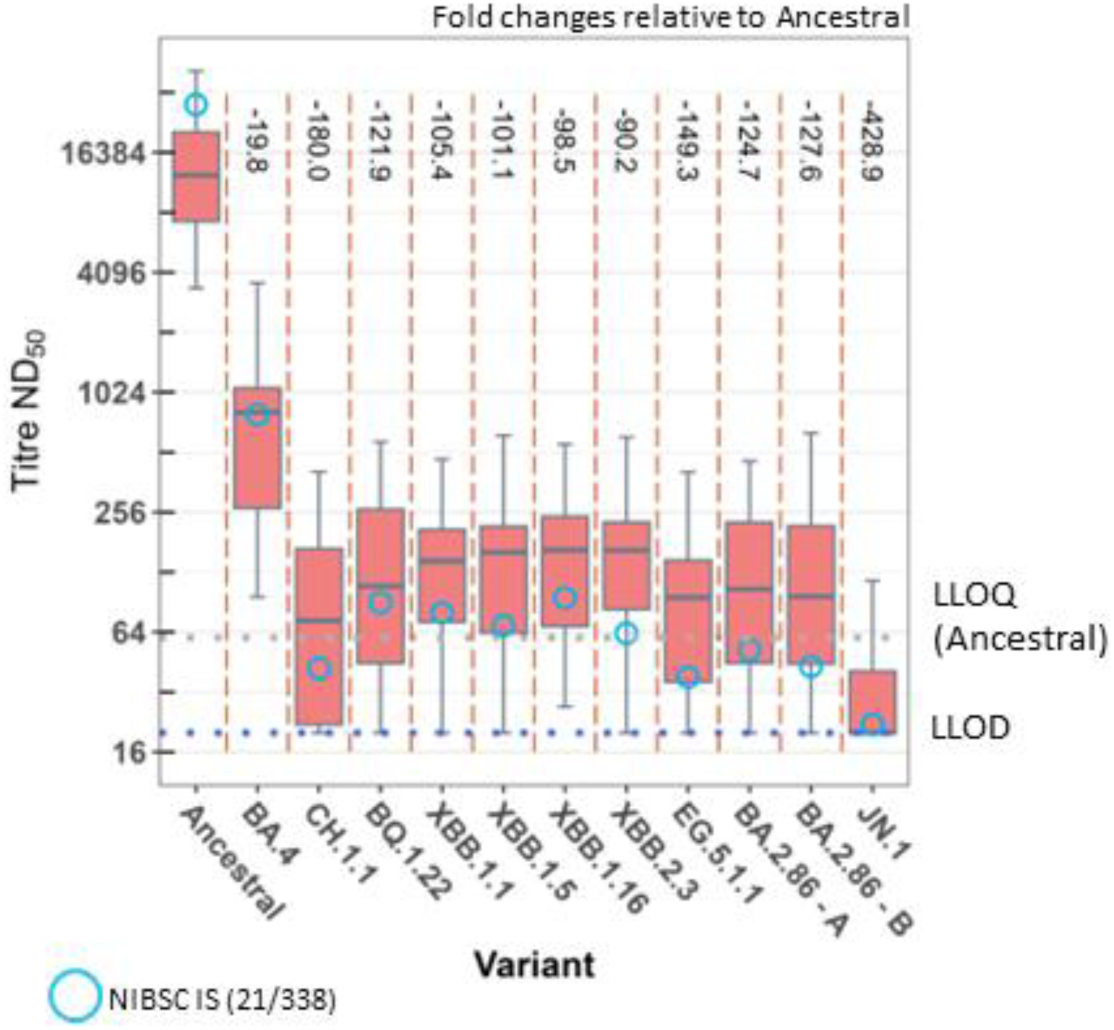
Neutralising antibody titres of a panel of sera/plasma from human volunteers who received three doses of an ancestral based vaccine as well as a BA.1 bivalent booster as determined by Focus Reduction Neutralisation Test (FRNT) against authentic SARS-CoV-2 Variants. Sera/plasma from participants who received four doses of a vaccine were assessed by FRNT with 11 authentic SARS-CoV-2 variants. The panel consists of sera (n=14) and plasma (n=8) samples from human participants who received three doses of an ancestral based vaccine as well as a BA.1 bivalent booster. Of these, 19 out of 22 had at least one confirmed breakthrough infection with SARS-CoV-2. Data are presented as median and interquartile ranges of titre (ND_50_). Fold-changes relative to ancestral virus were calculated by regression analysis using a linear mixed effect model and are presented where the difference is statistically significant (p<0.05, two-way ANOVA with Tukey’s *post hoc* test). The Lower Limit of Detection (LLOD) of 1/20 is indicated by the blue dotted line. The Lower Limit of Quantification (LLOQ) of 1/58 as determined for the ancestral assay, is indicated by the grey dotted line. Neutralisation titres of the NIBSC WR 21/338 is presented as a blue circle.

The international standard 21/338 was able to effectively neutralise all previous variants but only yielded a neutralisation titre of 22 against JN.1, which is nearing the LLOD (20). In addition, 12 of the 22 four-dose panel samples gave titres at or below the LLOD when tested against JN.1. The numerical breakdown of these data and the model estimates of geometric means are presented in Table S2.

Antigenic cartography is a powerful method for visualising the evolution of a pathogen by plotting the neutralisation titres of different panels of sera against successive variants or strains. It was originally developed as a means of monitoring changes in seasonal influenza virus and has since been used by numerous groups to compare SARS-CoV-2 variants [24–26]. Here, we use this technique to visualise the antigenic relationships of the SARS-CoV-2 variants tested against the panels of human sera or plasma described in this and our previous publication (Figure 3). The data plotted here include all titres from a panel of pre-Alpha convalescent plasma, triple-vaccinated individuals, boosted (BA.1/ancestral bivalent) individuals and the three standards (20/136, 21/334, 21/338). Each square of the map represents one antigenic unit which equates to a two-fold change in neutralisation titres. The ancestral virus is presented as a red circle and sits in the centre of the serum panels. Pre-Omicron variants were coloured as dark green circles and are situated within an inner circle around the serum panels. In the influenza field, vaccines are updated when circulating strains are at least two units away from the vaccine strain, thus marking it as genetically different [25]. Of the pre-Omicron variants, Beta and Zeta-FC are the furthest from ancestral virus with a gap of over 4 antigenic units, though in perpendicular directions from ancestral. Substantial diversity can be seen in post-Omicron variants which are grouped and coloured as Omicron-BA.1x (light green), Omicron-BA.2.x (sky blue), Omicron-BA.4/5.x (yellow), late Omicron (lilac), XE/XF recombinants (pink) and XBB recombinants (neon blue). However, some patterns can be seen, for example the recombinants XF and XE cluster with the virus from which their spike was derived. The recombinant XF is located with BA.1/BA.1.1 at approximately 5.5 units from ancestral and XE with BA.2/BA.2.12.1 at 4.5 units from ancestral. Also, the BA.4 and BA.5.2.1 cluster closely at approximately 5.5 units away from ancestral and, while the BA.4.6 sub lineage is a similar distance from ancestral, it is also approximately 4.5 units away from its parent, BA.4. The XBB family of variants cluster together at approximately 5.5 units from ancestral and within this cluster there are 1.2 antigenic units between the two most distant variants, XBB.1.16 and EG.5.1.1. The late Omicron variants (BQ.1.22, CH.1.1 and BA.2.86) form another separate cluster approximately seven antigenic units from ancestral with JN.1 approximately one antigenic unit further away from this cluster. Interestingly, BA.2.75.2 and BA.2.75.3 appear on the opposite side of the map to BA.2 with BA.2.75.2 located closer to the late Omicrons than to BA.2.75.3.

**Figure 3.**
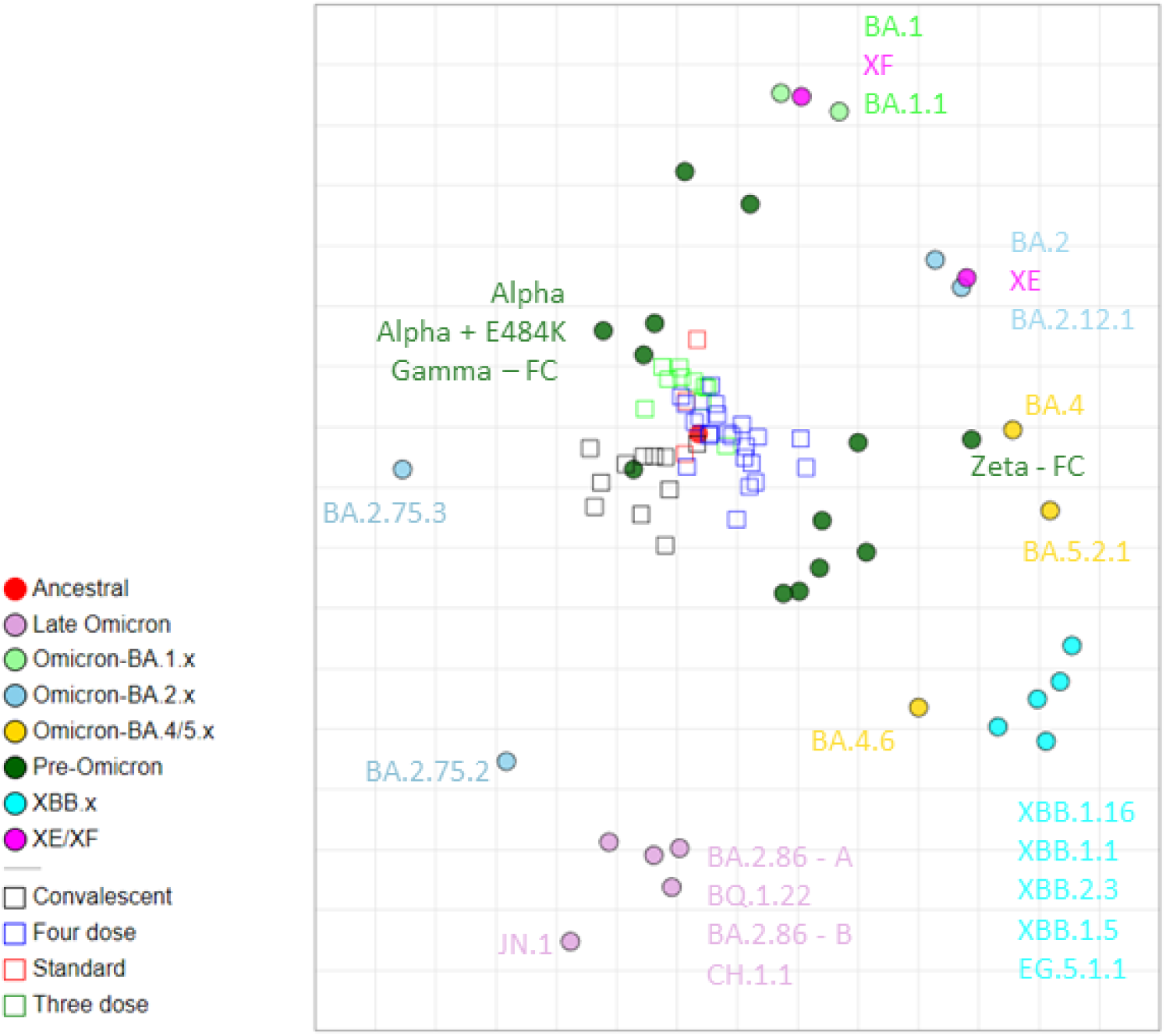
Antigenic map of SARS-CoV-2 variants assessed against human pre-Alpha convalescent, vaccinee sera or plasma and antibody standards. Median neutralisation titres (ND_50_) for human sera/plasma were generated against 34 authentic SARS-CoV-2 variants and plotted as antigenic maps. Serum/Plasma are plotted as open-coloured boxes and virus variants (antigens) as filled circles. Convalescent plasma (n=11) collected early in the pandemic (pre-Alpha), sera (n=10) from triple-vaccinated individuals and sera/plasma (n=22) from those who had received a BA.1 bivalent booster are shown as black, green, and blue boxes, respectively. The standards, WHO IS 20/136, NIBSC WR 21/334 and IS 21/338 are also plotted as red boxes. Virus variants are shown as coloured circles.

One of the known limitations of generating antigenic maps with human neutralisation data is the diversity of pre-existing immunity that is associated with multi-exposure sera (vaccine and naturally acquired). We therefore used single-exposure convalescent sera collected during our *in vivo* hamster infection studies to generate more antigenically defined maps. In Figure 4, we have plotted neutralisation titres of sera from hamsters collected on day 27 (ancestral and BA.1) or day 28 (BA.4, BA.5.2.1, XBB.1.5, XBB.1.16, XBB.2.3) after a single virus infection by the intranasal route. Each panel of sera was tested against the variant used as the challenge virus. Fold-changes against subsequent variants tested are presented as relative to the virus used for infection (Figure 4). Data for hamsters infected with ancestral, BA.1, BA.4 and BA.5.2.1 are described previously but also presented here [27, 28]. As our results from the human data indicated that the FCS substitution in the BA.2.86 – A virus did not affect neutralisation titres, repeat assays with these volume-limited sera were not performed with the second virus.

**Figure 4.**
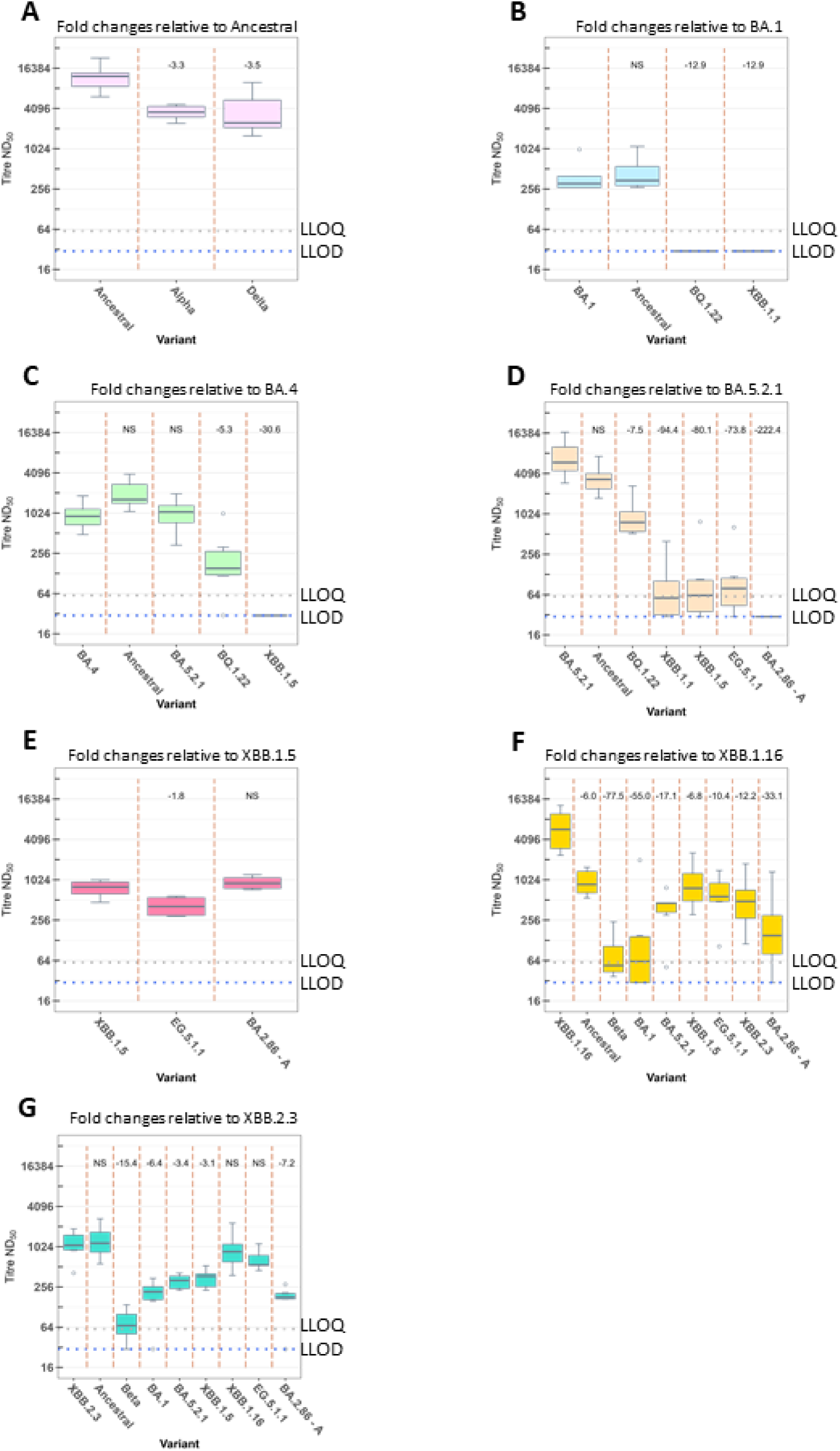
Neutralising antibody titres of panels of convalescent sera from hamsters experimentally infected with SARS-CoV-2 variants determined using an authentic virus Focus Reduction Neutralisation Test (FRNT) Sera was collected from Golden Syrian hamsters at day 27 or day 28 post-infection with ancestral (A), BA.1 (B), BA.4 (C), BA.5.2.1 (D), XBB.1.5 (E), XBB.1.16 (F) or XBB.2.3 (G) SARS-CoV-2 variants. Median neutralisation titres (ND_50_) were generated by FRNT using 14 authentic SARS-CoV-2 viruses. Data are presented as median and interquartile ranges of titre (ND_50_). Fold-changes relative to the homologous virus were calculated by regression analysis using a mixed effect model and are presented where the difference is statistically significant (p<0.05, two-way ANOVA with Tukey’s *post hoc* test). The Lower Limit of Detection (LLOD) of 1/30 is indicated by the blue dotted line. The Lower Limit of Quantification (LLOQ) of 1/58 as determined for the ancestral assay, is indicated by the grey dotted line. NS = not significant at the p<0.05 level.

Briefly, sera taken from hamsters infected with ancestral virus gave significantly lower GM titres with a 3.3- and 3.5-fold reduction for Alpha and Delta relative to ancestral, respectively. Sera taken from BA.1 convalescent hamsters showed a similar ability to neutralise both BA.1 and ancestral but were unable to neutralise BQ.1.22 and XBB.1.1 at the lowest dilution used here (1/30). For BA.4 convalescent hamster sera, BQ.1.22 and XBB.1.5 were the only variants which were significantly more resistant to neutralisation by these sera, with the latter resulting in GM titres all below the LLOD. In hamsters that were convalescent for BA.5.2.1, GM titres against ancestral virus relative to BA.5.2.1 were not significantly different. Titres against all other variants tested against BA.5.2.1 convalescent sera were significantly lower relative to BA.5.2.1. Against BQ.1.22, XBB.1.1, XBB.1.5 and EG.5.1.1 titres were above the LLOD meaning fold changes could be determined accurately with 7.5-, 94.4-, 80.1-, 73.8-fold reductions, respectively. Notably, when tested against BA.2.86 – A, none of the sera from BA.5.2.1 infected hamsters were able to neutralise the virus with all titres falling below the LLOD. GM titres of convalescent sera from hamsters infected with XBB.1.5 tested against EG.5.1.1 were significantly reduced relative to XBB.1.5 with a 1.8-fold reduction. When these sera were tested against BA.2.86 there was no significant difference in GM neutralising titres. Convalescent sera from hamsters experimentally infected with XBB.1.16 were tested against ancestral virus, Beta, BA.1, BA.5.2.1, XBB.1.5, EG.5.1.1, XBB.2.3 and BA.2.86 – A. GM titres for this panel against all variants were significantly reduced relative to XBB.1.16 with fold-reductions of 6.0, 77.5, 55.0, 17.1, 6.8, 10.4, 12.2 and 33.1, respectively. Finally, convalescent sera from hamsters experimentally infected with XBB.2.3 were able to neutralise ancestral, XBB.1.16 and EG.5.1.1 equally well to XBB.2.3. Against all other variants (Beta, BA.1, BA.5.2.1, XBB.1.5, BA.2.86 – A), the neutralisation titres for these sera were significantly lower with Beta displaying the most resistance to neutralisation by this panel. The numerical breakdown of these data and the model estimates of geometric means are presented in Tables S3-S9.

When these data were used to generate an antigenic map (Figure 5) the points corresponding to individual sera were more widely distributed compared to the human map. As expected, each antigenic variant clusters with sera from animals infected with homologous virus. Of the pre-Omicron variants, Beta is again the most antigenically distant at approximately three units from ancestral. BQ.1.22 is located in the same area of the map as Beta but another antigenic unit further from ancestral. The most antigenically distinct variant with the hamster map was XBB.1.1 at 5.5 units from ancestral and three units away from the nearest variant on the map, BA.1. The most contemporary variant tested here, BA.2.86 – A, is approximately five units from ancestral but clusters closely to XBB.1.5 with just over one antigenic units’ distance.

**Figure 5.**
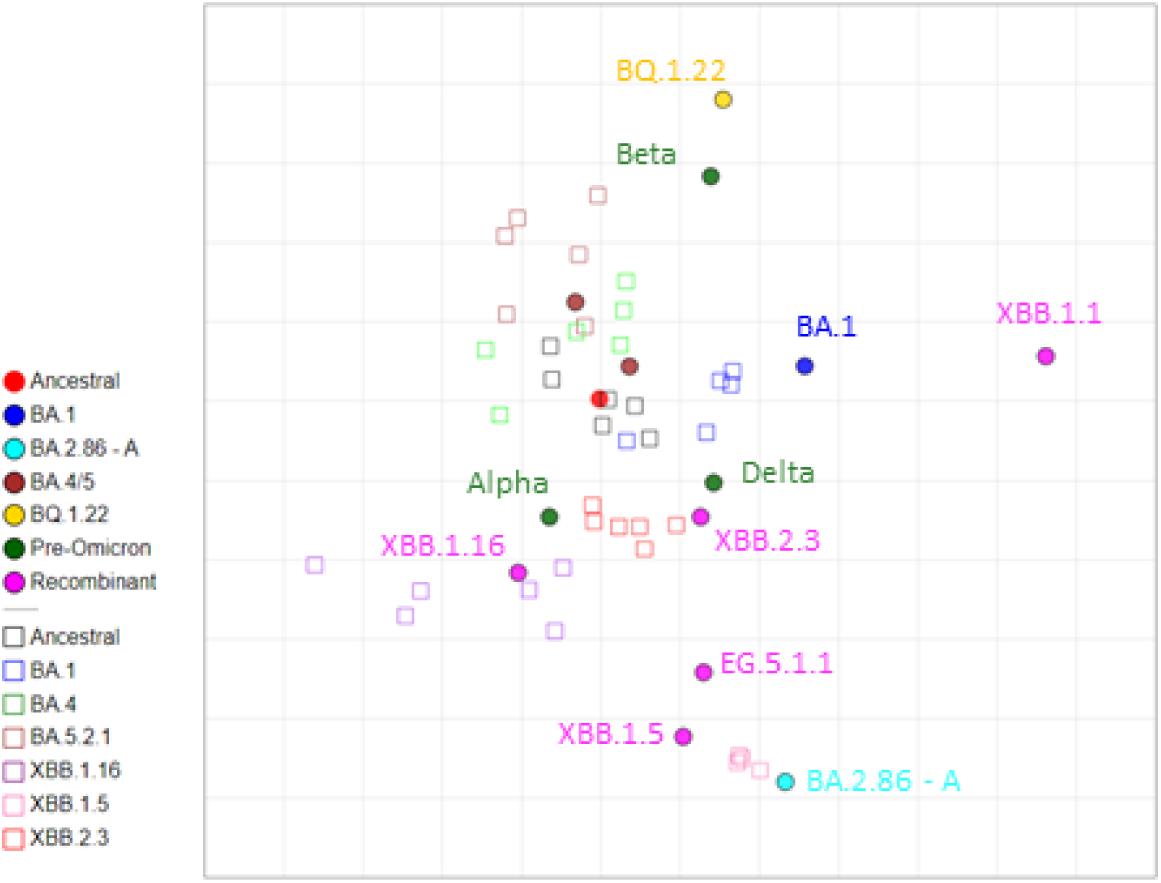
Antigenic map of SARS-CoV-2 variants assessed against convalescent sera collected from experimentally infected hamsters. Sera was collected from Golden Syrian hamsters at day 27 or day 28 post-infection with ancestral, BA.1, BA.4, BA.5.2.1, XBB.1.5, XBB.1.16 or XBB.2.3 SARS-CoV-2 variants. Median neutralisation titres (ND_50_) were generated by Focus Reduction Neutralisation Test (FRNT) using 14 authentic SARS-CoV-2 viruses and plotted using antigenic cartography. Serum/Plasma are plotted as open-coloured boxes and virus variants (antigens) as filled circles.

## Discussion

SARS-CoV-2 continues to cause worldwide disruption, despite the downgrading of the COVID-19 pandemic marking the end of the public health emergency. Since its emergence, SARS-CoV-2 has been acquiring fitness-enhancing mutations in response to changing population immunity driven by both infection and vaccination [29]. It is thought that additional pressures on the virus such as the clinical use of antiviral drugs, long-term infections in immunocompromised individuals and spillback events from zoonotic infections are also likely to contribute to the increased diversity in viral sequences [30, 31]. Here, we present our updated assessment of emerging SARS-CoV-2 variants in an authentic virus neutralisation assay against panels of sera or plasma from individuals who have received three doses of a COVID-19 vaccine as well as those who received a bivalent (BA.1/ancestral) booster. We also assessed neutralisation titres from experimentally infected hamsters that were convalescent for seven SARS-CoV-2 variants (ancestral, BA.1, BA.4, BA.5.2.1, XBB.1.5, XBB.1.16, XBB.2.3) and map both these and the human data antigenically.

In our previous publication we described a distinct difference in neutralisation titres against two different isolates of the Zeta variant when tested against our pre-Alpha convalescent panel. Another group from the University of Geneva showed a similar large reduction in neutralisation titres with an early pandemic convalescent panel of sera against Zeta that was not seen against triple vaccinated individuals [32]. We therefore tested our two Zeta isolates against our three-dose vaccinee panel, however these viruses continued to show substantial resistance to neutralisation by this panel with a significant difference between the two isolates. A sequence comparison with data kindly supplied by the University of Geneva revealed identical spike sequences for all three viruses but with a three amino acid deletion in the N gene of the virus discussed in the Bekliz paper (208-210). Zeta – BEI and the University of Geneva virus also have three additional non-synonymous mutations in Orf1a leading to amino acid changes: V1071A, P1810L and S3149F. With no differences in the main viral proteins exposed to neutralising antibodies (spike, membrane, envelope), it appears that the sequence is not responsible for the variation in titres seen here. Other potential explanations were described in our previous paper and include the potential impact of secretion of free spike protein, defective interfering particles and differences in spike glycosylation [12, 33–37].

Our data from individuals who had received three doses of an ancestral spike-based vaccine indicated that antibodies against the two variants, BA.2.75.2 and BA.4.6, were less able to neutralise these viruses with titres nearing the LLOD. BA.2.75.2 has three additional amino acid changes relative to BA.2.75.3 (identical spike to BA.2.75) which are thought to increase immune evasion (S:R346T, S:F486S, S:D1199N) [38, 39]. Similarly, BA.4.6 also contains S:R346T relative to its parent, BA.4. Our neutralisation data supports this prediction with a higher immune escape of BA.2.75.2 compared to BA.4.6 and agrees with others who show similar trends [40, 41].

Further testing of this population demonstrated that the sera in this panel had no detectable neutralising activity against the later variants, BQ.1.22 and XBB.1.1. The BQ.1.22 variant used in this study has the same spike sequence as the BQ.1.1 variant, which was dominant in December 2022, with three additional changes relative to its parent BA.5 (S:R346T, S:K444T, S:N460K). Similarly, the recombinant XBB.1.1, circulating around the same time, had changes in several of the same locations including S:R346T, S:L368I, S:N460K, S:F486S, S:R493Q and S:F490S. These two variants were also resistant to neutralisation by panels of triple vaccinated volunteers in other studies as well as in those with breakthrough infections with BA.1 and BA.2 [42, 43].

In the UK, as boosters containing bivalent antigens were deployed in the Autumn of 2022, we continued our assessment of emerging variants with samples from individuals who had received a BA.1/ancestral booster. The majority also had at least one breakthrough infection between February 2021 and December 2022 (Alpha to XBB.1.5). Results from this panel showed that neutralisation titres for the two variants which were no longer neutralised by the three-dose panel (BQ.1.22 and XBB.1.1) were effectively neutralised by this four-dose panel. Indeed, all the subsequent recombinants and Omicron sub-lineages tested here showed similar neutralisation titres for all variants between CH.1.1 to BA.2.86. This includes the two BA.2.86 isolates tested here, with and without the presence of an amino acid change in the FCS (S:R682W), with no significant difference seen for sera tested against the two viruses. These data are in alignment with others who have demonstrated that changes in the FCS are not important in relation to virus neutralisation *in vitro* but alter how the virus behaves *in vivo* [19–22]. Following the emergence of the BA.2.86 sub-lineage, JN.1 (August 2023) and its subsequent declaration by the WHO as a VOI (19^th^ December 2023), we also isolated and investigated this variant [5]. JN.1 has a further amino acid change in the RBD in addition to BA.2.86 (S:L455S) which has generated concern due to changes at this position in other variants being associated with increased escape from humoral immunity [44].

All variants tested displayed increased resistance to neutralisation to this four-dose panel relative to both ancestral and BA.4, with similar GM titres for all except JN.1 (CH.1.1, BQ.1.22, XBB.1.1, XBB.1.5, XBB.1.16, XBB.2.3, EG.5.1.1, BA.2.86). This aligns with results presented by other groups using both pseudovirus (PSV) and authentic virus assays against panels of vaccinated (three- and four-dose) with additional break-through infection in a globally diverse population [45–52]. It is also of interest that the GM titres, up to and including BA.2.86, are comparable despite the genetic diversity in the variants tested against our four-dose panels. This is likely due to the range of variants to which these individuals have acquired natural immunity via break-through infections. The fold-reduction of our neutralisation titres against JN.1 relative to BA.2.86 are comparable to two studies using PSV assays which demonstrated an approximate two-fold reduction [53, 54]. Two other studies, however, demonstrated similar titres against both BA.2.86 and JN.1 [55, 56]. The serum samples used in these studies were collected more recently than our own panel which could explain the differences seen with our own results. A common observation from all of the published data is that sera from those with a breakthrough infection since the emergence of late XBB variants such as XBB.1.5 or EG.5.1 have higher titres and broader cross-protection between variants. Another study generated neutralisation data for the JN.1 variant used sera from XBB.1.5 vaccinated individuals. In that study, the neutralisation titres were consistently higher against all variants, including JN.1, post vaccination. BA.2.86 was not investigated in this study, however the fold-reductions for JN.1 relative to XBB.1.5 (2.9 to 4.3-fold) were comparable to those estimated by our own study for these variants (4.2-fold). We note however that, due to vaccination campaigns targeting at-risk subgroups within the population, the sera or plasma from individuals in our panel of boosted vaccinees are part of a limited subset of the population considered to be at risk. Whilst this is informative in terms of the effects of vaccines on immunity it may be less representative of levels of vaccine-induced immunity in the general population.

Antigenic mapping of the human data shows co-localisation of the pre-Omicron variants in the centre of the map with a larger antigenic spread following the emergence of Omicron and recombinants. This is comparable to maps presented by others using human data, though direct comparisons are challenging given the diversity in vaccination, circulating variant exposure and breakthrough infections in different panels of human sera used between studies [24, 57–60]. For example, many will have had different vaccination schedules/ formulations and exposure to unidentified viral variants in addition to human genetic diversity leading to a range of humoral immune responses. However, while distances between variants in different maps may vary, the general pattern is the same with pre-Omicron variants localised to the centre and clusters of related variants moving out from the centre in a chronological manner. As others have shown, the XBB family of variants cluster further away from ancestral than BA.4/5 and late Omicron variants such as BQ.1.22, CH.1.1 and BA.2.86 are more distant again [52, 61]. Another study which mapped JN.1 antigenically also showed this variant as distinct from other variants, but it did not have data from other late Omicron variants to enable comparison of how these may group together [62].

One limitation of ours and others human antigenic maps is the clustering of sera in one location on the map, corresponding to the ancestral spike that drove the pandemic and was used as the basis of the first-generation vaccines. This effect is less prominent when using the neutralisation titres from sera taken from hamsters 27- or 28-days post-infection with a single, known SARS-CoV-2 variant at a standard infectious dose. Antigenic mapping of these data highlighted two antigenically distinct viruses: BQ.1.22 and XBB.1.1. Although the most contemporary variant to date, BA.2.86, is one of the furthest antigenically from ancestral, it clustered closely with XBB.1.5. This suggests that antibodies raised against the XBB.1.5 spike antigen have good cross-reactivity against BA.2.86, thus supporting the transition to XBB.1.5 based vaccines. Others have also shown XBB.1.1, BQ.1.1 and BA.2.86 to be distinct in antigenic maps generated with hamster or mouse data [52, 61]. Using single exposure sera from hamsters has the added benefits of being generated by a known priming antigen, given at the same dose with the duration since infection being known and fixed. These are therefore more predictive of antigenic relationships when mapped. Our hamster data also indicate that a BA.5 spike alone does not generate a detectable neutralising antibody response to one of the most recent variants, BA.2.86. Taken together, these data suggested a turning point in the virus’ evolution and signalled to vaccine developers that an updated vaccine was required. On 18 May 2023 the WHO Technical Advisory Group on COVID-19 Vaccine Composition (TAG-CO-VAC) published their recommendations to develop vaccines to target contemporary XBB-descendent lineages [63].

Numbers of sequenced SARS-CoV-2 genomes are decreasing due to reduced global surveillance activity since the end of the pandemic. As a result, identifying variants of concern in a timely manner is increasingly challenging. This was highlighted by the apparent sudden emergence of BA.2.86 in August 2023 despite some models retrospectively predicting it had been circulating since May 2023 [48, 64]. Continued monitoring of emerging variants and their assessment in programmes such as Agility is essential to build on existing knowledge of immune evasion of contemporary and newly emerging variants and to better advise policymakers and vaccine manufacturers on future strategies.

## Methods

### Human vaccinee sera and antibody standards

SARS-CoV-2 variants were tested against two panels of vaccinee samples collected from staff volunteers. The first panel, consisting of sera from individuals who had received three doses of an ancestral spike-based COVID-19 vaccine, was collected as part of the ESCAPE study as previously described [12, 65]. The second panel was assembled from staff members at MHRA (n=8 plasma samples) and UKHSA (n=14 sera samples) who had received a fourth vaccination consisting of bivalent (BA.1/ancestral) vaccine. Samples were collected at a median of 78 days after the fourth vaccination. When tested for the presence of anti-N antibodies, 19 of the 22 individuals had evidence of convalescence for naturally acquired SARS-CoV-2 infection. Dates of previous infections were provided by 14 individuals which identified an infection range of February 2021 to December 2022. Two reference materials were included in this study: a working reagent for anti-SARS-CoV-2 immunoglobulin (NIBSC 21/234), which is a pool of several pre-Alpha convalescent plasma, and the International Standard for antibodies to SARS-CoV-2 VOC (NIBSC 21/338), which is a pool of 265 individuals who are both vaccinated and convalescent following Alpha, Beta or Delta infection [66, 67].

### Virus bank preparation and quality control checks

Unless otherwise stated in Table 1, viruses were isolated from sequence-confirmed clinical swabs at UKHSA. SARS-CoV-2 variants were isolated and propagated on Vero/hSLAM cells (ECACC 04091501) before being subjected to quality control checks as previously described [12]. Briefly, titrations were performed using a focus-forming assay with immunostaining for nucleocapsid to visualise foci [68]. Sighting for each variant into the Focus Reduction Neutralisation Test (FRNT) to calculate dilution at which a median of 130 foci could be counted in non-neutralised control wells was performed. Whole genome sequencing (SISPA-Illumina) was performed on final virus assay banks, as described previously, to confirm presence of FCS S1/S2 boundary, check for mixed bases as well as confirm variant identity [12, 69–71]. A 12-nucleotide insertion (after position 21608) and nine-nucleotide deletion (21633-21641) in BA.2.86 results in short illumina reads in this region which map poorly to the reference sequence NC_045512.2. Therefore, BA.2.86, illumina sequence was mapped to Oxford Nanopore Technologies (ONT) consensus sequence in place of NC_045512.2. Sequence data of the virus banks used in the neutralisation tests in this study are available in Supplementary Data File S1.

### FRNT and statistics

Median neutralising antibody titres (ND_50_) were measured with FRNT and calculated with a Probit regression analysis using R v4.1.2 (R Core Team, Vienna, Austria) as described previously [11, 12, 68]. Log_10_-transformed ND_50_ titres were analysed using a mixed-effects linear model in R, as described previously [12, 72].

### Antigenic cartography

Antigenic maps were generated using the Racmacs (https://acorg.github.io/Racmacs/) package in R as described previously [24, 73]. The number of dimensions was set to 2, number of optimisations to 500 and minimum column basis to none. Neutralisation titres from human and hamster sera were used to generate two separate maps.

### Hamster sera

All *in vivo* experimental work was conducted under the authority of a UK Home Office approved project licence that has been subject to local ethical review at either UK Health Security Agency Porton Down or Medicines and Healthcare products Regulatory Agency, South Mimms by their respective Animal Welfare and Ethical Review Body (AWERB) as required by the *Home Office Animals (Scientific Procedures) Act 1986*.

Sera collected at day 27 post-infection from hamsters exposed to ancestral virus (n=6) and BA.1 (n=5) or day 28 post-infection with BA.4 (n=6) and BA.5.2.1 (n=6). These hamster sera were generated as previously described [27, 28]. Sera was collected from hamsters infected with XBB1.16 and XBB.2.3 (n=6 per variant) with a target dose of 1×10^4^ FFU/200 µL via intranasal instillation (method previously described [27, 28]). Blood samples were collected during necropsy at 28 days post infection.

Sera from XBB.1.5 convalescent hamsters were collected as follows. Four hamsters (approximately 10 weeks old) were administered 1×10^5^ FFU/50µL intranasally of XBB.1.5. After infection, oral swabs were taken to confirm infection by lateral flow and RT-qPCR. Hamsters were terminated at 28 days post-infection and clotted whole blood was centrifuged (1,000RPM, 5 mins) to obtain serum.

## Acknowledgements

The authors gratefully acknowledge the support from the Porton Sequencing Laboratory and the Biological Services Porton at the UK Health Security Agency, Porton Down, United Kingdom. The authors thank our staff volunteers for providing serum and plasma samples so that we can assemble our testing panels. We also acknowledge Samuel Wilks and Derek Smith for their help and advice with using the Racmacs package. The authors are grateful to Professor Isabella Eckerle and Dr Meriem Bekliz for sharing sequence data for their Zeta variant and useful discussions.

## Ethics

The ESCAPE study was conducted in accordance with the Declaration of Helsinki and the Governance Arrangements for Research Ethics Committees (GAfREC). It was approved by the PHE Research Ethics and Governance Group (R&D REGG Ref NR0190, March 2020). Informed consent was obtained from all subjects involved. Donations from MHRA staff members were collected with written informed consent from donors, study was approved by the MHRA local ethics committee (HuMAC ref: 20/05/SK).

