## Supplementary Tables 1-9 for "Evaluation of the cross reactivity of neutralising antibody response in vaccinated human and convalescent hamster sera against SARS-CoV-2 variants up to and including JN.1 using an authentic virus neutralisation assay"

| **Table S1** – Summary of geometric mean neutralisation titres from a focus-reduction assay (FRNT) for a panel of 10 triple vaccinated human sera against SARS-CoV-2 variants and their fold-changes relative to ancestral virus. | | | |
| --- | --- | --- | --- |
| **Variant** | **Geometric mean of titres (ND_50­_)** | **Fold-change relative to ancestral virus** | **95% confidence intervals** |
| Ancestral | 5230.0 | NA | 3440.0 - 7930.0 |
| Zeta - FC | 199.0 | -26.2* | 136.0 - 291.0 |
| Zeta - BEI | 671.0 | -7.8* | 504.0 - 892.0 |
| BA.1 | 182.0 | -28.7* | 97.4 - 340.0 |
| BA.1.1 | 144.0 | -36.4* | 93.4 - 221.0 |
| BA.2 | 199.0 | -26.2* | 142.0 - 280.0 |
| BA.2.12.1 | 166.0 | -31.4* | 115.0 - 241.0 |
| BA.2.75.3 | 168.0 | -31.0* | 96.3 - 295.0 |
| BA.2.75.2 | 46.1 | -113.3* | 25.2 - 84.6 |
| BA.4 | 104.0 | -50.2* | 71.1 - 152.0 |
| BA.4.6 | 49.8 | -105.0* | 30.5 - 81.1 |
| BA.5.2.1 | 65.4 | -79.9* | 43.2 - 98.8 |
| BQ.1.22 | 26.5 | -196.9* | 15.8 - 44.6 |
| XE | 162.0 | -32.3* | 114.0 - 229.0 |
| XF | 151.0 | -34.5* | 89.5 - 256.0 |
| XBB.1.1 | 20.5 | -255.5* | 19.4 - 21.5 |
| Significant difference (p<0.05), as determined by a two-way ANOVA with Tukey’s HSD *post hoc* test are indicated by *. | | | |

| **Table S2** – Summary of geometric mean neutralisation titres from a focus-reduction assay (FRNT) for a panel of 22 human sera/plasma taken from those who had received three ancestral based vaccines as well as a BA.1/ancestral bivalent booster against SARS-CoV-2 variants. Fold-changes relative to ancestral virus are shown. | | | |
| --- | --- | --- | --- |
| **Variant** | **Geometric mean of titres (ND_50­_)** | **Fold-change relative to ancestral virus** | **95% confidence intervals** |
| Ancestral | 12400.0 | NA | 8970.0 - 17100.0 |
| BA.4 | 624.0 | -19.8* | 393.0 - 992.0 |
| CH.1.1 | 68.7 | -180.0* | 43.7 - 108.0 |
| BQ.1.22 | 102.0 | -121.9* | 62.6 - 165.0 |
| XBB.1.1 | 117.0 | -105.4* | 79.4 - 174.0 |
| XBB.1.5 | 122.0 | -101.1* | 80.2 - 187.0 |
| XBB.1.16 | 126.0 | -98.5* | 85.0 - 186.0 |
| XBB.2.3 | 137.0 | -90.2* | 96.9 - 194.0 |
| EG.5.1.1 | 82.9 | -149.3* | 54.7 - 126.0 |
| BA.2.86 - A | 99.3 | -124.7* | 65.6 - 150.0 |
| BA.2.86 - B | 97.0 | -127.6* | 63.0 - 149.0 |
| JN.1 | 28.9 | -428.9 | 22.8 - 36.6 |
| Significant difference (p<0.05), as determined by a two-way ANOVA with Tukey’s HSD *post hoc* test are indicated by *. | | | |

| **Table S3** – summary of geometric mean neutralisation titres from a focus-reductions assay (FRNT) for a panel of sera taken from Syrian hamsters (n=6) at day 27 post a single challenge with ancestral SARS-CoV-2. | | | |
| --- | --- | --- | --- |
| **Variant** | **Geometric mean of titres (ND_50­_)** | **Fold-change relative to ancestral virus** | **95% confidence intervals** |
| Ancestral | 11600 | NA | 7130.0 - 19000.0 |
| Alpha | 3560 | -3.3* | 2720.0 - 4650.0 |
| Delta | 3350 | -3.5* | 1550.0 - 7250.0 |
| Significant difference (p<0.05), as determined by a two-way ANOVA with Tukey’s HSD *post hoc* test are indicated by *. | | | |

| **Table S4** – summary of geometric mean neutralisation titres from a focus-reductions assay (FRNT) for a panel of sera taken from Syrian hamsters (n=5) at day 28 post a single challenge with the BA.1 SARS-CoV-2 variant. | | | |
| --- | --- | --- | --- |
| **Variant** | **Geometric mean of titres (ND_50­_)** | **Fold-change relative to BA.1** | **95% confidence intervals** |
| BA.1 | 388 | NA | 196.0 - 769.0 |
| Ancestral | 440 | 1.1 | 212.0 - 914.0 |
| BQ.1.22 | 30 | -12.9* | 30.0 - 30.0 |
| XBB.1.1 | 30 | -12.9* | 30.0 - 30.0 |
| Significant difference (p<0.05), as determined by a two-way ANOVA with Tukey’s HSD *post hoc* test are indicated by *. | | | |

| **Table S5** – summary of geometric mean neutralisation titres from a focus-reductions assay (FRNT) for a panel of sera taken from Syrian hamsters (n=6) at day 28 post a single challenge with the BA.4 SARS-CoV-2 variant. | | | |
| --- | --- | --- | --- |
| **Variant** | **Geometric mean of titres (ND_50­_)** | **Fold-change relative to BA.4** | **95% confidence intervals** |
| BA.4 | 917 | NA | 558.0 - 1510.0 |
| Ancestral | 1920 | 2.1 | 1130.0 - 3260.0 |
| BA.5.2.1 | 943 | 1.0 | 494.0 - 1800.0 |
| BQ.1.22 | 172 | -5.3* | 50.9 - 583.0 |
| XBB.1.5 | 30 | -30.6* | 30.0 - 30.0 |
| Significant difference (p<0.05), as determined by a two-way ANOVA with Tukey’s HSD *post hoc* test are indicated by *. | | | |

| **Table S6** – summary of geometric mean neutralisation titres from a focus-reductions assay (FRNT) for a panel of sera taken from Syrian hamsters (n=6) at day 28 post a single challenge with the BA.5.2.1 SARS-CoV-2 variant. | | | |
| --- | --- | --- | --- |
| **Variant** | **Geometric mean of titres (ND_50­_)** | **Fold-change relative to BA.5.2.1** | **95% confidence intervals** |
| BA.5.2.1 | 6670.0 | NA | 3380.0 - 13200.0 |
| Ancestral | 3330.0 | -2.0 | 1960.0 - 5640.0 |
| BQ.1.22 | 892.0 | -7.5* | 465.0 - 1710.0 |
| XBB.1.1 | 70.7 | -94.4* | 24.7 - 202.0 |
| XBB.1.5 | 83.3 | -80.1* | 23.2 - 299.0 |
| EG.5.1.1 | 90.4 | -73.8* | 28.7 - 285.0 |
| BA.2.86 – A | 30.0 | -222.4* | 30.0 - 30.0 |
| Significant difference (p<0.05), as determined by a two-way ANOVA with Tukey’s HSD *post hoc* test are indicated by *. | | | |

| **Table S7** – summary of geometric mean neutralisation titres from a focus-reductions assay (FRNT) for a panel of sera taken from Syrian hamsters (n=4) at day 28 post a single challenge with the XBB.1.5 SARS-CoV-2 variant. | | | |
| --- | --- | --- | --- |
| **Variant** | **Geometric mean of titres (ND_50­_)** | **Fold-change relative to XBB.1.5** | **95% confidence intervals** |
| XBB.1.5 | 745 | NA | 430.0 - 1290.0 |
| EG.5.1.1 | 410 | -1.8* | 229.0 - 734.0 |
| BA.2.86 – A | 923 | 1.2 | 625.0 - 1360.0 |
| Significant difference (p<0.05), as determined by a two-way ANOVA with Tukey’s HSD *post hoc* test are indicated by *. | | | |

| **Table S8** – summary of geometric mean neutralisation titres from a focus-reductions assay (FRNT) for a panel of sera taken from Syrian hamsters (n=6) at day 28 post a single challenge with the XBB.1.16 SARS-CoV-2 variant. | | | |
| --- | --- | --- | --- |
| **Variant** | **Geometric mean of titres (ND_50­_)** | **Fold-change relative to XBB.1.16** | **95% confidence intervals** |
| XBB.1.16 | 5550.0 | NA | 2580.0 - 12000.0 |
| Ancestral | 922.0 | -6.0* | 577.0 - 1470.0 |
| Beta | 71.7 | -77.5* | 33.0 - 156.0 |
| BA.1 | 101.0 | -55.0* | 17.9 - 569.0 |
| BA.5.2.1 | 325.0 | -17.1* | 119.0 - 885.0 |
| XBB.1.5 | 819.0 | -6.8* | 363.0 - 1840.0 |
| EG.5.1.1 | 536.0 | -10.4* | 208.0 - 1380.0 |
| XBB.2.3 | 454.0 | -12.2* | 166.0 - 1240.0 |
| BA.2.86 - A | 168.0 | -33.1* | 41.9 - 672.0 |
| Significant difference (p<0.05), as determined by a two-way ANOVA with Tukey’s HSD *post hoc* test are indicated by *. | | | |

| **Table S9** – summary of geometric mean neutralisation titres from a focus-reductions assay (FRNT) for a panel of sera taken from Syrian hamsters (n=6) at day 28 post a single challenge with the XBB.2.3 SARS-CoV-2 variant. | | | |
| --- | --- | --- | --- |
| **Variant** | **Geometric mean of titres (ND_50­_)** | **Fold-change relative to XBB.2.3** | **95% confidence intervals** |
| XBB.2.3 | 1050 | NA | 594.0 - 1850.0 |
| Ancestral | 1200 | 1.1 | 659.0 - 2170.0 |
| Beta | 68 | -15.4* | 38.0 - 122.0 |
| BA.1 | 165 | -6.4* | 65.6 - 413.0 |
| BA.5.2.1 | 306 | -3.4* | 234.0 - 400.0 |
| XBB.1.5 | 341 | -3.1* | 241.0 - 482.0 |
| XBB.1.16 | 866 | -1.2 | 451.0 - 1660.0 |
| EG.5.1.1 | 639 | -1.6 | 444.0 - 919.0 |
| BA.2.86 - A | 146 | -7.2* | 63.2 - 336.0 |
| Significant difference (p<0.05), as determined by a two-way ANOVA with Tukey’s HSD *post hoc* test are indicated by *. | | | |
